# The Mitochondrial Genome and Epigenome of the Golden Lion Tamarin from Fecal DNA using Nanopore Adaptive Sequencing

**DOI:** 10.1101/2021.05.27.446055

**Authors:** Nicole Wanner, Peter A. Larsen, Adam McLain, Christopher Faulk

**Affiliations:** Department of Animal Sciences, University of Minnesota, College of Food, Agricultural, and Natural Resource Sciences, Saint Paul, MN, USA; Department of Veterinary and Biomedical Sciences, College of Veterinary Medicine, University of Minnesota, Saint Paul, MN, USA; Department of Biology and Chemistry, College of Arts and Sciences, SUNY Polytechnic Institute, Utica, NY, USA

**Author notes:** **Corresponding author:** Christopher Faulk, Assistant Professor, Department of Animal Sciences, College of Food, Agricultural, and Natural Resource Sciences, University of Minnesota, 1988 Fitch Ave., Saint Paul, MN 55108. **Contributing authors** Nicole Wanner Peter Larsen Adam T. McLain.

**Keywords:** Mitochondria, DNA methylation, DNA hydroxymethylation, Poop, Primates, Lion Tamarin

## Abstract

The golden lion tamarin (*Leontopithecus rosalia*) is an endangered Platyrrhine primate endemic to the Atlantic coastal forests of Brazil. Despite ongoing conservation efforts, genetic data on this species remains scarce. Complicating factors include limitations on sample collection and a lack of high-quality reference sequences. Here, we used nanopore adaptive sampling to resequence the *L. rosalia* mitogenome from feces, a sample which can be collected non-invasively. Adaptive sampling doubled the fraction of both host-derived and mitochondrial sequences compared to sequencing without enrichment. 258x coverage of the *L. rosalia* mitogenome was achieved in a single flow cell by targeting the unfinished genome of the distantly related emperor tamarin (*Saguinus imperator*) and the mitogenome of the closely related black lion tamarin (*Leontopithecus chrysopygus*). The *L. rosalia* mitogenome has a length of 16,597 bp, sharing 99.68% sequence identity with the *L. chrysopygus* mitogenome. A total of 38 SNPs between them were identified, with the majority being found in the non-coding D-loop region. DNA methylation and hydroxymethylation were directly detected using a neural network model applied to the raw signal from the MinION sequencer. In contrast to prior reports, DNA methylation was negligible in mitochondria in both CpG and non-CpG contexts. Surprisingly, a quarter of the 642 CpG sites exhibited DNA hydroxymethylation greater than 1% and 44 sites were above 5%, with concentration in the 3’ side of several coding regions. Overall, we report a robust new mitogenome assembly for *L. rosalia* and direct detection of cytosine base modifications in all contexts.

## Introduction

The golden lion tamarin (*Leontopithecus rosalia*) is an endangered Platyrrhine primate endemic to the Atlantic coastal forests of Brazil (Rylands et al. 2002). It is a member of the family Callitrichidae, a taxonomic grouping that includes marmosets, tamarins and lion tamarins (Groves 2001). Although once teetering on the brink of extinction, golden lion tamarins have benefitted from a successful captive breeding and reintroduction program that has seen their numbers climb from a few hundred individuals in the wild to several thousand. In addition to wild individuals, several hundred animals are maintained in captivity globally as part of the captive breeding and reintroduction program, and stud books are maintained to promote genetic diversity in these animals and avoid inbreeding (Kierulff, M. C. M.; Ruiz-Miranda, C. R.; de Oliveira, P. Procópio; Beck, B. B.; Martins, A.; Dietz, J. M.; Rambaldi, D. M.; Baker 2012). Golden lion tamarins are gregarious, living in social groups typically centered around a monogamous breeding pair and their dependent infants, juveniles and subadults (Coimbra-Filho and Mittermeier 1973). They are omnivorous, feeding on a wide array of plants and animals including flowers, fruits, and small vertebrates and invertebrates. Golden lion tamarins also play an important ecological role as seed dispersers (Lapenta, MJ; Procópio-de-Oliveira, P.; Kierulff, MCM; Motta-Junior 2008). Genetic data from *L. rosalia* remains scant, however. Increased availability of genetic data will benefit breeding and conservation efforts for this species.

Mitochondria are the powerhouse of the cell and contain their own genome (Goldsmith et al. 2020). The mitogenome in animals is small, circular, and mutates more rapidly than the nuclear genome. Two characteristics make it especially useful for phylogenetic comparisons and non-invasive sampling. First, it is present in many more copies per cell than the nuclear genome, making it easier to recover from degraded samples such as feces or ancient DNA. Second, its higher mutation rate enables more accurate delineation of closely related species. The first mitochondrial sequences of *L. rosalia* were published in 2008 and 2011 and were limited to the sequences of cytochrome b and the hypervariable region of the displacement control region (D-loop) (Perez-Sweeney et al. 2008; Matauschek et al. 2011). A more complete version of the mitogenome became available in 2013, but it contained gaps and shared surprisingly limited homology with the closely related black lion tamarin (*L. chrysopygus*) (Finstermeier et al. 2013). To create a more accurate comparison of the *L. rosalia* mitogenome to its sister species, we chose to resequence its mitochondrial genome to high coverage using a novel technique with easily collected fecal samples.

Nanopore sequencing is based on electrical signals generated by cylindrical protein pores as nucleic acids pass through (Lu et al. 2016). The MinION sequencer from Oxford Nanopore Technologies (ONT) is field-portable with minimal reagent requirements. It can provide long reads with an average read length of 20kb and occasional single reads over 1 Mb in length, limited only by the molecular weight of the input DNA. Since mitochondria have ∼16kb genomes, read lengths are sufficient to cover the entire mitogenome. However, in fecal samples, the mixture of DNA sources presents a challenge to bulk sequencing.

Traditionally, targeted sequencing of specific loci is performed by enzymatic or PCR enrichment prior to sequencing (Gilpatrick et al. 2020). Recently, a form of computational enrichment called adaptive sampling has been developed (Payne et al. 2020b). This method allows for bulk sequencing of whole genomic DNA combined with locus-specific enrichment by rejecting off target reads. Crucially, adaptive sampling can enrich target regions up to 30-fold, dependent on sequence length and percentage of the genome, enough to bring target loci to high enough coverage for analysis. Here, we employed adaptive sampling in order to obtain sufficient coverage of the mitogenome for accurate assembly.

From an epigenetic perspective, mitochondrial DNA methylation has been a matter of debate (Sharma et al. 2019b). Determining its methylation pattern has been challenging due to unique characteristics including resistance to bisulfite transformation, which is used in nearly all epigenetic methods (Chandler et al. 2017). Direct sequencing of native genomic DNA preserves base modifications which can then be detected with nanopore sequencing. Neural network models are capable of providing simultaneous calls of nucleic acid sequence and base modifications from unenriched genomic DNA input (Wick et al. 2019). Due to the recent availability of models for calling DNA hydroxymethylation, we are the first to report native detection at base-pair resolution in mitochondria. The combination of adaptive sampling, long-reads, and epigenetic information, drove our selection of the Oxford Nanopore MinION to resequence the mitogenome of *L. rosalia*.

Fecal DNA serves as a rich source of host information that can be collected non-invasively from wild populations and processed for DNA extraction in the field (Wang et al. 2018). Fecal microbiomes from nanopore sequencing have been the subject of multiple studies (Cuscó et al. 2019; Moss et al. 2020; Shanmuganandam et al. 2019), however, host DNA enrichment by nanopore sequencing from fecal samples has not yet been documented. Due to degradation of DNA by digestive processes, choosing high abundance targets such as mitochondria naturally increases their sequencing frequency at the cost of sequence read length.

We describe an improved assembly of the mitogenome of the golden lion tamarin extracted from a fecal sample. We find that this species is most closely related to the black lion tamarin based on the high level of sequence identity. This finding is consistent with prior taxonomic studies using genetics and morphology, which found the two species to be closely related (Perez-Sweeney et al. 2008; Mundy and Kelly 2001; Rosenberger and Coimbra-Filho 2008). We confirm the pattern of diverging mutations falls mostly within the non-coding D-loop. Our method resulted in 285x coverage from a single flow cell, allowing high confidence in the consensus. Sequence polishing, error-correcting, and assembly resulted in a circular contig of 16,597 bp in length. We directly measured DNA methylation and hydroxymethylation levels from the nanopore signal level read data using the Megalodon tool provided by ONT. We find that DNA methylation is negligible across the entire mitogenome, in line with human mtDNA (Goldsmith et al. 2020). Surprisingly, we find elevated levels of DNA hydroxymethylation only in the CpG context, suggesting biological function of this mark in mitochondria. We conclude that fecal samples provide a rich source of host DNA suitable for nanopore sequencing, providing a new capability for field use in conservation research.

## Results

### Bulk Fecal DNA yields a large quantity of highly fragmented DNA

Wild-collected feces represent an abundant supply of host genomic, metabolomic, and metagenomic information. We collected a fresh, wet fecal sample from a captive golden lion tamarin at the Utica Zoo. Two kit-based methods of DNA extraction were used with manufacturer protocols designed to enrich the host DNA fraction and yielded similar results. A total of 9 μg of DNA was derived from 400 mg of fecal material and used for library prep and nanopore sequencing. Library prep with 2 μg input yielded between 200 and 700 ng of product per reaction. Mean read length from untargeted nanopore sequencing was 1230 bp, indicating short fragment length likely due to digestive degradation.

### Nanopore adaptive sampling enriches host DNA from feces

A nanopore MinION flowcell was loaded with 500 ng of DNA and sequenced for 24 hours. Since over 99% of fecal DNA is derived from microbiome and digesta, enriching the small fraction of host-derived fragments makes better use of limited sequencing capacity. We used adaptive sampling to enrich for sequences that matched to the unfinished genome of a distantly related species, *Saguinus imperator* (emperor tamarin). This target genome consists of 3.4 Gb total sequence divided into 1,666,189 scaffolds. *S. imperator* was chosen over the more closely related *Leontopithecus chrysopygus* (black lion tamarin) since the latter did not have an available nuclear genome.

Our initial sequencing run yielded 15 million reads containing 6.3 Gb of sequence with a mean read length of 411 bp, indicating that adaptive sampling was rejecting most reads in less than 1 second since the read rate averaged over 400 bases/s. We identified 0.8% of reads matching *S. imperator*, derived from the fecal sample of *L. rosalia* DNA, compared to 0.39% without enrichment, therefore adaptive sampling was successful in enriching host tamarin sequences (Table 1).

**Table 1.**
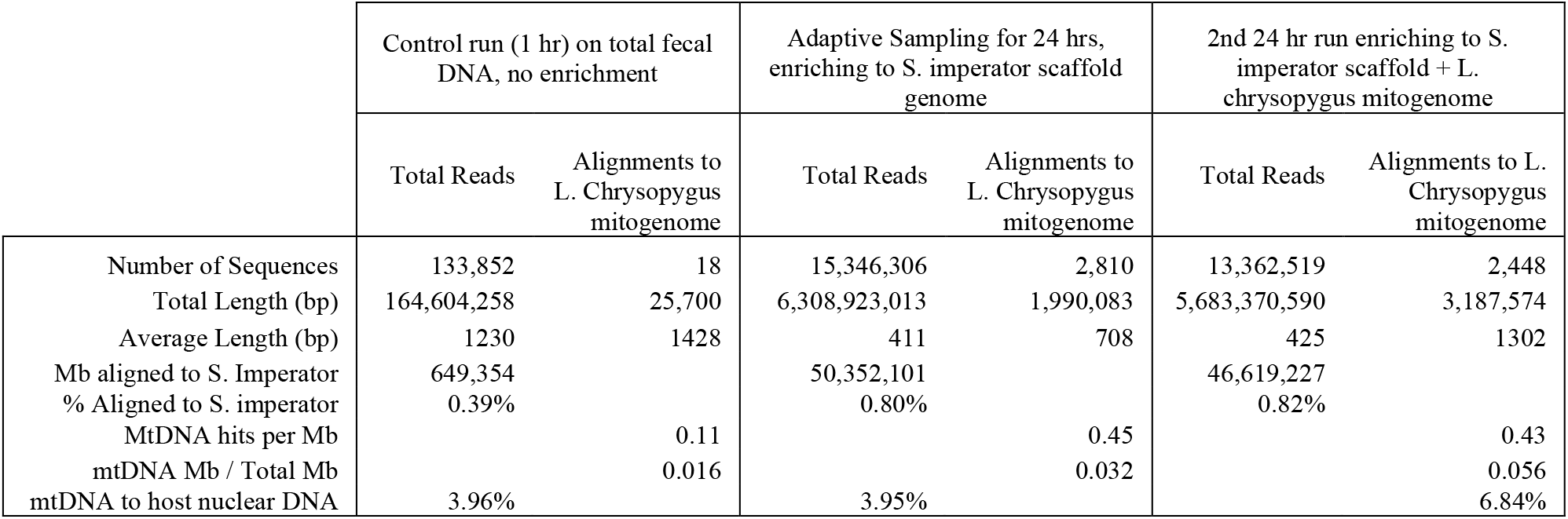
Mapping statistics for Control and Adaptive Sampling Runs

|  | Control run (1 hr) on total fecal DNA, no enrichment |  | Adaptive Sampling for 24 hrs, enriching to *S. imperator* scaffold genome |  | 2nd 24 hr run enriching to *S. imperator* scaffold + *L. chrysopygus* mitogenome |  |
| --- | --- | --- | --- | --- | --- | --- |
|  | Total Reads | Alignments to *L. Chrysopygus* mitogenome | Total Reads | Alignments to *L. Chrysopygus* mitogenome | Total Reads | Alignments to *L. Chrysopygus* mitogenome |
| Number of Sequences | 133,852 | 18 | 15,346,306 | 2,810 | 13,362,519 | 2,448 |
| Total Length (bp) | 164,604,258 | 25,700 | 6,308,923,013 | 1,990,083 | 5,683,370,590 | 3,187,574 |
| Average Length (bp) | 1230 | 1428 | 411 | 708 | 425 | 1302 |
| Mb aligned to *S. Imperator* | 649,354 |  | 50,352,101 |  | 46,619,227 |  |
| % Aligned to *S. imperator* | 0.39% |  | 0.80% |  | 0.82% |  |
| MtDNA hits per Mb |  | 0.11 |  | 0.45 |  | 0.43 |
| mtDNA Mb / Total Mb |  | 0.016 |  | 0.032 |  | 0.056 |
| mtDNA to host nuclear DNA | 3.96% |  | 3.95% |  |  | 6.84% |

### Mapping mitochondrial reads

To identify mitochondrially derived reads, we initially mapped the total read set to the previously reported *L. rosalia* reference mitogenome (NC_021952). We found 4585 matching reads with a mean length of 1104 bp. However, the alignment revealed a high number of SNPs, gaps, 3’ artifacts, and low fidelity to the existing reference. Our resulting contig had 96.83% identity across only 92% of the query length to *L. rosalia* NC_021952, quite unexpectedly divergent for members of the same species. An unusually large number of reads were mapped to a small, disconnected fragment on the 3’ end of this reference (Figure S1). To determine whether these were sequencing artifacts or errors in the reference, we aligned our reads to the closely related *L. chrysopygus* mitogenome (accession NC_037878)(de Freitas et al. 2018) which yielded fewer reads (2810) of shorter average length (708 bp) but much more uniform coverage. Our resulting contig had 99.86% identity over >99% of the length of this mitogenome without gaps, suggesting it would provide a better reference for further analyses.

### Mitochondrial DNA is enriched by adaptive sampling

To determine whether adaptive sampling also enriched mitochondrial DNA along with nuclear DNA, we compared reads from a non-adaptively sampled control run vs. the first 24 hours of an adaptive sampling run targeting the *S. imperator* scaffold genome. We found a 4-fold increase in number of mtDNA sequences per Mb when mapping to the *L. chrysopygus* mitogenome from the *S. imperator* enriched reads (Table 1). By read percentage, host mtDNA doubled from 0.016% to 0.032% of the total fecal DNA, precisely mirroring the doubling seen in total host genomic DNA enrichment. There was no difference in the mitochondrial vs. host nuclear DNA ratio with adaptive sampling, with mtDNA making up 3.95% of the host DNA in both. This result was expected since we enriched for an entire tamarin genome, not just mtDNA specifically.

### Higher quality targets increase enriched read length but not sampling efficiency

Next, we explored whether adaptive sampling efficiency could be improved by adding a more accurate target for enrichment. The *S. imperator* genome initially used for adaptive sampling contains fragments covering the entire *S. imperator* mitogenome but with very poor quality. When assembled and compared against the high-quality *L. chrysopygus* mitogenome, the two share only ∼70% sequence identity, indicating many errors. In contrast, *L. chrysopygu*s matches 95% to the existing *L. rosalia* mtDNA (NC_021952) and >99% to the *L. rosalia* mitogenome that we ultimately assembled. We reasoned that inclusion of the *L. chrysopygus* mitogenome along with the *S. imperator* full genome might improve adaptive sampling efficiency since it would share much higher sequence identity with *L. rosalia* mtDNA. A second aliquot of the same library was run on the MinION for an additional 24 hours, with both the *S. imperator* scaffold genome and *L. chrysopygus* mitogenome as enrichment targets, resulting in 5.6 Gb of additional reads.

In the second run, reads matching the *L. chrysopygus* mitogenome were 1302 bp in length, double the 708 bp read length of initial run. This improved length did not result in greater overall enrichment, however; enrichment remained at a 4-fold increase over background. Increased read length did result in 60% greater coverage of the mitogenome, improving contig coverage from 106x in the first run to 179x in the second run. Read length improvement and coverage is illustrated in figure 1. The region of lowest coverage was the D-loop, while the nearby large subunit rRNA had the highest average coverage. Interestingly, in the second run the ratio of mtDNA to host nuclear DNA nearly doubled from 3.95% to 6.83% due to the increased read length from having a more accurate mitogenome target.

**Figure 1.**
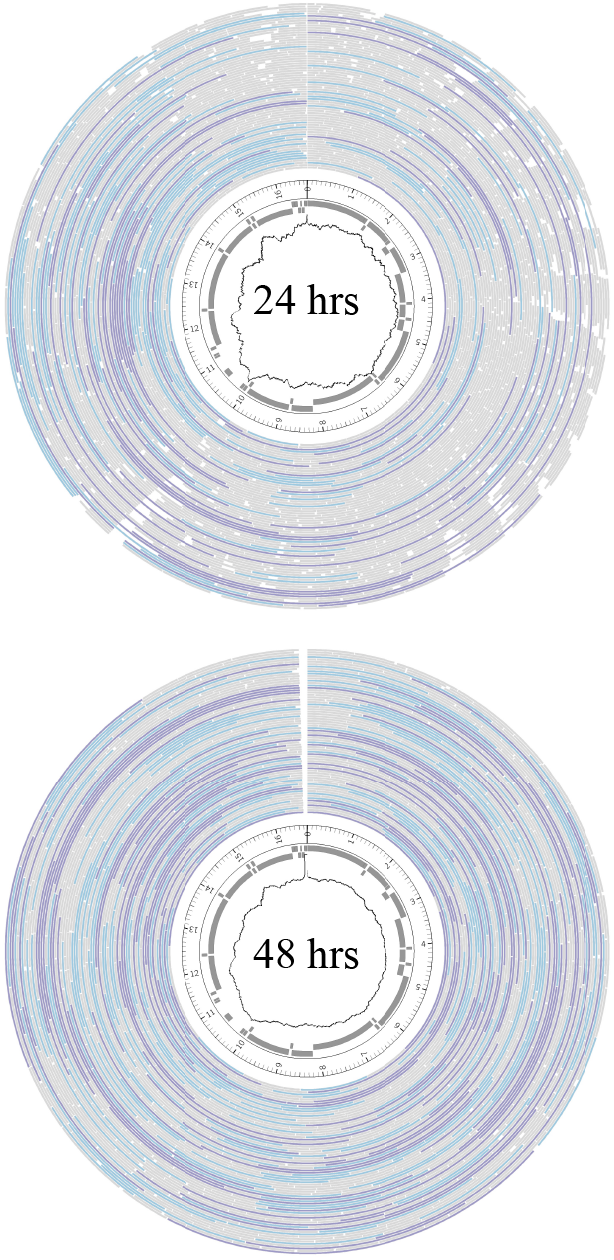
Coverage density of the mitogenome. Outer layer shows mapped reads where >2kb are purple, 1-2kb are blue, <1kb are gray. Inner layers show gene position and coverage. For the first 24 hour run (top), the minimum coverage is 52x, at the D-loop, maximum = 168x at the large subunit-rRNA. For the combined data over 48 hours (bottom), minimum coverage is 169x at the D-loops, and maximum = 392x at the large subunit-rRNA.

### Nanopore adaptive sampling tolerates large target enrichment sequence divergence

Both rounds of adaptive sampling resulted in 4-fold improvement in mitogenome coverage vs. non-targeted sequencing, in line with Oxford Nanopore’s guidance. This indicates that adaptive sampling can tolerate at least 30% sequence identity divergence from the target while still enriching as highly as compared to a perfectly matching target. For subsequent analyses, we combined reads from both runs.

### Mapping reveals high coverage of host mitochondrial DNA

Overall, there were 5516 reads aligning to *L. chrysopygus* mtDNA with a mean length of 984 bp, resulting in 285x read coverage with improved continuity particularly at the 3’ end (Fig. S1). Assembly of reads with Flye yielded a circular draft contig of 16,592 bp with 99.86% identity to *L. chrysopygus* over 99% of its length. This draft assembly contained several indels proximal to homopolymeric regions as seen in other reports, suggesting polishing could improve the sequence quality(Baeza 2020). When compared to the existing *L. rosalia* (NC_021952) reference this assembly matched at only 96.83% identity for 92% of its length, representing a strong improvement in both measures. The next nearest BLAST match in the NCBI nucleotide database was Goeldi’s marmoset (*Callimico goeldii*) at 85.18% identity with 97% query length coverage.

### Polishing of the assembly

Identification of open reading frames was performed with MITOS2 and revealed that the indels in the draft assembly were causing multiple frameshift mutations. Both the Oxford Nanopore tool Medaka and the 3^rd^ party tool Nanopolish were used for polishing with the former outperforming the latter. Medaka eliminated all frameshifts. Comparison of the draft and polished assemblies revealed 5 indels all proximal to homopolymeric regions and 1 SNP, with identity to *L. chrysopygus* dropping the match to 99.63% for >99% of its length. This is a substantial improvement compared to the existing *L. rosalia* reference which matched to the polished contig at 95.95% identity for 95% of its length. MITOS2 annotation of the polished assembly found that all frameshift mutations were resolved, and no manual intervention was required. The polished assembly was rearranged to place *COX1* at the start position and submitted to NCBI under accession number MZ262294.

### Mitogenome organization is highly conserved with mutations preferentially in the D-loop

The mitogenome of *L. rosalia* has a total length of 16,597 bp (Fig. 2). Our assembly is shorter than the previous reference by 275 bp, and is very similar in size to the closely related *L. chrysopygus* (16,618 bp) (de Freitas et al. 2018). Coding regions are nearly identical to *L. chrysopygus* with 2 amino acid changes in *NADH* and none in any other. Gene order is the same as in humans and other primates. We then investigated mutations that accrued since the divergence of *L. rosalia* and *L. chrysopygus*. A total of 38 SNPs are mapped on figure 2. As expected, the majority of SNPs are concentrated within the non-coding D-loop. Only 16 SNPs are located throughout the rest of the genome.

**Figure 2:**
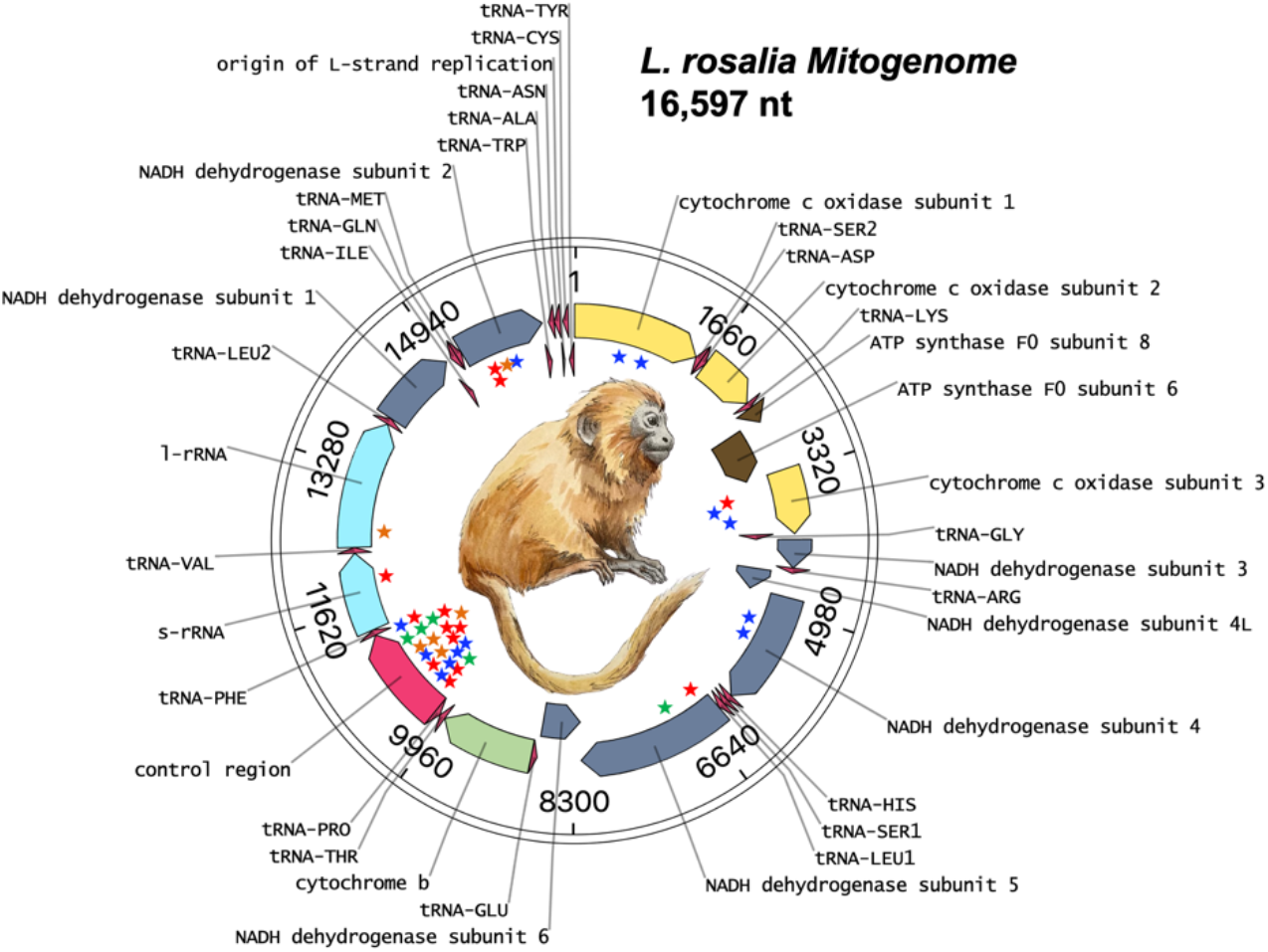
The mitogenome of the Golden Lion Tamarin. The *L. rosalia* mitogenome is 16,597 bp in length. Stars represent SNPs in comparison to the closely related Black Lion Tamarin, *L. chrysopygus*. The majority of SNPs are in the D-Loop control region.

### DNA methylation is negligible in mitochondria at CpG and CH sites

Native genomic DNA contains modifications of interest in epigenetic analyses. We chose a sequencing library preparation kit that excludes PCR steps in order to preserve these modifications for downstream analyses. As others have reported both the presence and absence of DNA methylation in mitochondria using a variety of methods, we sought to resolve this controversy by leveraging the ONT’s neural network models to call modified bases. We used Megalodon software with neural network models trained to detect 5’methylation (5mC) and 5’hydroxymethylation (5hmC) at cytosines in any context (Table 2). The first and second 24-hour runs were analyzed separately as technical replicates (Fig 3).

**Table 2.** DNA modification summary

|  | Total Sites | >1% methylation |  |  | >5% methylation |  |  | Coverage |  |
| --- | --- | --- | --- | --- | --- | --- | --- | --- | --- |
|  |  | 1st run | 2nd run | # replicated | 1st run | 2nd run | # replicated | 1st run | 2nd run |
| 5mC CpG | 642 | 11 | 13 | 0 | 2 | 2 | 0 | 43x | 63x |
| 5hmC CpG | 642 | 328 | 309 | 196 | 124 | 67 | 44 | 43x | 63x |
| 5mC CH | 5469 | 0 | 0 | 0 | 0 | 0 | 0 | 49x | 73x |
| 5hmC CH | 5469 | 183 | 189 | 18 | 4 | 0 | 0 | 49x | 73x |

**Figure 3.**
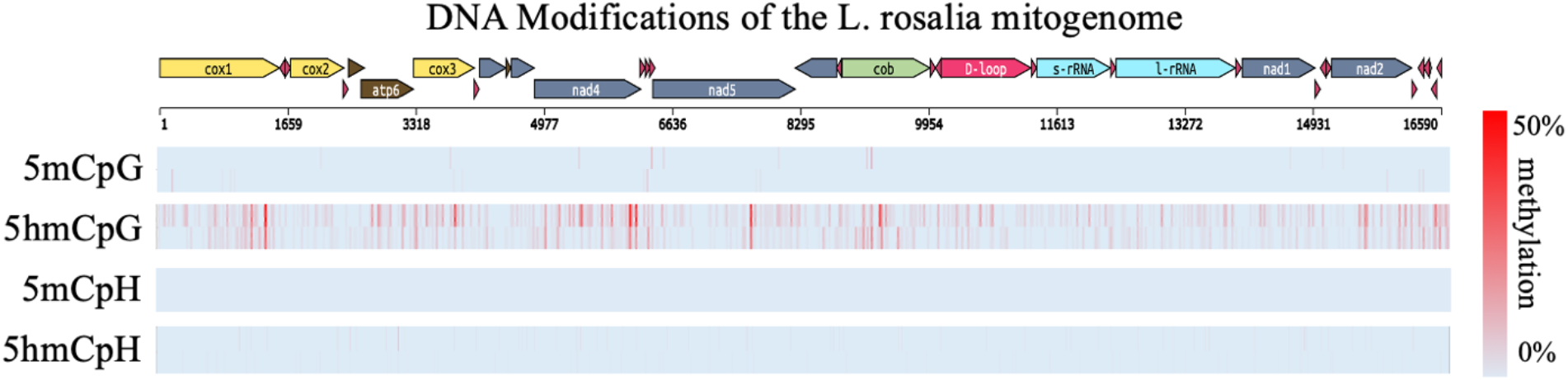
DNA modifications of the L. rosalia mitogenome. Cytosine modifications are shown mapped to the mitogenome of L. rosalia. The modifications are illustrated with replicates above and below from the first and second runs. Hydroxymethylation in the CpG context has the highest prevalence and replication. Percentage of total modified base reads are shown from blue to red.

Across CpG sites within the mitogenome, DNA methylation of was seen in just 13 of the 642 total sites at a greater 1% level, and none of these had greater than 0% in the technical replicate. Only 2 sites were detected when a cutoff of >5% methylation was set, and neither was above 0% in the technical replicate. In contrast, hydroxymethylation greater than 1% was called at nearly half the 642 sites, with 196 of these replicating between runs. When filtering for CpG sites with hydroxymethylation greater than 5%, 44 sites were detected and replicated between runs. At 4 sites 5hmC exceeded 20% in both replicates. Interestingly, 5hmC is concentrated on the 3’ side of several genes including *COX1, COX3, NAD4*, and *NAD5* indicating a non-uniform distribution and likely biological function. We found no significant correlation in methylation between strands at any threshold.

In non-CpG context, identified by the dinucleotide ambiguity code CH where H stands for “any nucleotide except G”, no cytosines were methylated in any of the 5469 possible CH sites. For hydroxymethylation, we found 3.4% of sites with detectible levels above 1%, however, only 18 of these sites replicated. At the 5% threshold, no replication remained. Taken together these findings indicated strong support for the presence of hydroxymethylation in the CpG context, with neither methylation nor hydroxymethylation present outside of CpG sites. (Supplementary File 1).

## Discussion

The purpose of this study was to selectively enrich and sequence host DNA from a non-invasively collected sample using nanopore adaptive sequencing, assemble a mitogenome, and assess for epigenetic modifications. We chose the golden lion tamarin, *L. rosalia*, as it represents an endangered primate with an unsequenced genome and a mitochondrial genome in need of improvement. The fraction of host DNA sequenced from feces was less than 1% of the total reads in line with other studies,(Sharma et al. 2019a) but this fraction was doubled with the use of adaptive sampling. Remarkably, our coverage of the mitogenome was high enough to accurately assemble a full length contig. These findings suggest that mitogenomes make the best target for nanopore sequencing from fecal DNA.

Recently, the mitogenome of the related Brazilian buffy-tufted-ear marmoset (*Callithrix aurita*) was assembled by genomic skimming with a nanopore sequencer (Malukiewicz et al. 2021). Genomic skimming uses low coverage sequencing and leverages naturally high copy number sequences (e.g. repeats, and mitogenomes) to assemble high coverage of those regions. It had high concordance with a Sanger sequenced replicate, however it resulted in only 9x coverage of the mitogenome and required two full flow cells as opposed to our 285x coverage with a single flow cell. Closer to our target species, de Freitas et al. recently used short-read sequencing to generate a high quality mitogenome of *L. chrysopygus* with over 3000x coverage, providing us with an accurate reference for our nanopore adaptive sampling method (de Freitas et al. 2018). That mitogenome required bootstrapping alignments with manual k-mer adjustments, whereas our long-read pipeline assembled an error-free mitogenome despite lower coverage. Both methods leverage the ability to use related species as alignment targets.

The enrichment of mitochondrial DNA was surprising despite its overabundance compared to nuclear chromosomes. Generally, the ratio of mtDNA to nuclear DNA is about 0.1%, despite containing hundreds to thousands more copies per cell in most somatic tissues (Robin and Wong 1988). Here we found a 40-fold increase in this ratio, at 3.96% mtDNA to nuclear DNA. By including an accurate copy of the *L. chrysopygus* mitogenome, we increased that ratio to 6.84%, doubled the length of aligned reads, and quadrupled the hits per megabase of reads. Our increased efficiency may be a product of the sample source. Bulk stool contains highly fragmented digested DNA, and the described method may not hold for less degraded mitochondrial DNA. Nanopores draw in only linearized DNA, therefore any mtDNA molecules must be degraded enough to lose their native circular conformation before sequencing. Even though we did not use any double strand break treatments prior to sequencing (e.g. enzymatic treatment, sonication, hydrodynamic shearing), we still sequenced 285-fold coverage of the mitogenome, as compared to less than 0.1x of the nuclear genome.

Oxford Nanopore guidance indicates that longer targets and read lengths are correlated with higher enrichment. Our lower level of enrichment (4-fold) over non-enriched sequencing is likely due to short fragment lengths. Indeed, when sequencing human mitochondrial DNA from high molecular weight libraries from liver cells, Goldsmith et al. found reads averaging over 80% of the entire mitogenome (Goldsmith et al. 2020). Meanwhile, adaptive sampling yields best results with targets greater than 15 kb and read lengths greater than 10kb (Payne et al. 2020b). A second reason for our low level of enrichment was the poor match of the emperor tamarin mtDNA sequences. With the better *L. chrysopygus* match our aligned reads lengthened to 1300 bp, slightly longer than the average read length of the control run with no rejection taking place. This result indicates that DNA fragmentation in the feces was the limiting factor.

The presence of mitochondrial pseudogenes in the nuclear genome did not appear to bias our results, given the strong concordance of our assembly to the *L. chrysopygus* mtDNA which was created with careful emphasis to eliminating nuclear coded mt-pseudogenes (de Freitas et al. 2018). The coding structure of our assembly mirrors the *L. chrysopygus*, with SNPs primarily in the D-loop. SNP accumulation in the D-loop was expected due to its non-coding status.

Our full sequence of the *L. rosalia* mitogenome shares 99.63% sequence identity with the published *L. chrysopygus* mitogenome, indicating very recent divergence of these sister species. Moreover, the assembly fits better with the known phylogenetic distances within Callitrichidae. The sister taxa relationship between these species had been under debate until 2001 when photoreceptor intron sequences unambiguously established their relationship (Mundy and Kelly 2001). In, 2008 the first mitochondrial sequences, of the D-loop, confirmed their close relationship (Perez-Sweeney et al. 2008). We extend this evidence by showing near perfect synteny throughout the coding regions.

The presence of mitochondrial DNA modifications has long been debated. Here we report essentially zero DNA methylation at every CpG position in the mitogenome in stark contrast to other reports of low but consistent methylation in mtDNA. Mitochondrial methylation using a nanopore sequencer has been reported by at least four groups (Aminuddin et al. 2020; Bicci et al. 2021; Lüth et al. 2021; Goldsmith et al. 2020). All of these studies found low but measurable mtDNA methylation, averaging less than 7% across a variety of conditions and cell types, though a few specific CpG sites had over 20%. These studies all used Nanopolish which does not distinguish 5mC from 5hmC. Since bisulfite-based methods also cannot distinguish 5mC from 5hmC and, they are similarly prone to overestimating the level of 5mC. It is also well-known that bisulfite does not transform circular DNA with high efficiency, again leading to an overestimate of 5mC (Mechta et al. 2017). Nanopore alleviates these concerns by avoiding bisulfite conversion, and all DNA molecules sequenced through a pore are linearized by necessity. Here we used Megalodon’s neural network model capable of detecting 5hmC to avoid Nanopolish’s 5mC bias. In support of our findings, a recent careful analysis of mitochondria methylation by Bicci et al. validated nanopore sequencing across multiple primary and cancer cell lines (Bicci et al. 2021). When accounting for all confounders, they found negligible DNA methylation as we have here. We suggest that previous findings of mitochondrial 5mC were artefacts of either bisulfite or miscalled nanopore basecalling.

We found very high levels of 5’hydroxymethylation within the mitogenome in the CpG context. Interestingly, the most well-cited work on mitochondrial base modifications by Shock et al. reported 10x greater levels of 5hmC than 5mC, using antibody-based methods (Shock et al. 2011). Biological function of 5’hydroxycytosine is strongly suggested for two reasons. First, we found no non-CpG modifications at all. This argues for reader proteins capable of recognizing cytosines in dinucleotide context rather than these modifications resulting from non-specific oxidation processes modifying cytosines by chance. Interestingly we saw no correlation in 5hmC between strands, indicating a non-palindromic oxidation process. This stranded-ness in mitochondrial base modification was seen by Dou et al., though they used bisulfite methods an called it as 5mC (Dou et al. 2019). Second, we see enrichment on the 3’ side of several genes, showing non-uniform distribution which is a hallmark of function (McLain and Faulk 2018). Taken together, our data suggests a specific biological function of 5hmCpG in mitochondria.

Epigenetic data is challenging to recover from wild species but has been done in hyenas and is expanding to other vertebrates (Laubach et al. 2019; De Paoli-Iseppi et al. 2017). Therefore, our study serves two purposes. First, it confirms the ability to detect base modifications by nanopore sequencing of mitogenomes (Chandler et al. 2017). Second, it provides proof-of-concept that epigenetic analyses can be performed on fecal derived samples, which is applicable to both host nuclear genome methylation and the fecal microbiome, even potentially including food species in digesta.

It is important to consider some caveats to our method. Nanopore accuracy is still low relative to short-read and Sanger sequencing, though neural network models are improving at a rapid pace (Wick et al. 2019). Polishing helps as indicated by Medaka’s ability to removing all frame shift inducing indels from our assembly. Modified base calling is also improving rapidly and is also dependent on models trained preferably for a specific modification and with taxa-specific data.

Interest in long-read sequencing of vertebrate mitogenomes continues to increase. A study by Formenti et al. recently sampled 100 species and corrected many long-standing errors in reference mitogenomes (Formenti et al. 2020). However, their approach is unsuitable for the most easily available tissue, fecal DNA, since the first step is to remove short fragments less than 10kb. Here we show the strength of short fragments and long-read technology together to generate accurate assemblies and epigenomes useful for comparative genomics (Colwell et al. 2018). Accurate genomic resources are critical to the conservation of endangered species like the Golden Lion Tamarin.

## Materials and methods

### Sample collection

A single fecal sample weighing 10 grams was collected from the environment of the two golden lion tamarins at the Utica Zoo between 2 and 8 hours post defecation. The sample was deposited by the female of a pair, Arie, who was 6 years old at the time of collection. All samples were collected under the approval of the Utica zoo. Fecal samples are exempt from IACUC protocol approval at the University of Minnesota.

### DNA extraction

Qiagen QIAamp DNA stool mini kit (cat no. 51504) was used with the protocol, “Isolate of DNA from Stool for Human DNA Analysis” supplied by manufacturer. This kit yielded approximately 4 μg of purified DNA from 200 mg of fecal sample. Additionally, the Omega Bio-tek E.Z.N.A. Stool DNA kit (cat no. D4015-00) was used also with 200 mg of stool and manufacturer’s protocol, “DNA Extraction and Purification from Stool for Human DNA Detection” and yielded a similar quantity of DNA.

Library prep was performed using the Oxford Nanopore Technologies (ONT) SQK-LSK109 “Genomic DNA by Ligation” protocol with the following changes: 1) NEBNext products were used for end-repair as suggested by ONT (cat no. M6630, E7546, and E6056), 2) Axygen AxyPrep Mag PCR Clean-up beads were substituted for Agencourt AMPure beads, 3) magnetic beads were optionally diluted to 25% of original volume with in-house prepared carboxy bead dilution buffer (https://bomb.bio/protocols/) for cost savings with no loss in recovery efficiency (Oberacker et al. 2019).

### Nanopore adaptive sequencing

Sequencing was conducted on a MinION sequencer on a FLO-min106 flow cell with pore chemistry R9.4 for two runs of 24 hrs each. Between runs the flow cell was washed with nuclease from the Flow Cell Wash Kit (WSH004) and loaded with storage buffer. Live basecalling was performed using the fast basecalling model in ONT basecalling software, Guppy v4.5.4, GPU enabled with a GeForce RTX 2080Ti. Adaptive sampling was enabled in MinKNOW core v.4.2.5 and set to enrich sequences matching to the *Saguinus imperator* (emperor tamarin) genome, SagImp_v1_BIUU (accession PRJNA399417), scaffold assembly. After sequencing, reads were basecalled again with Guppy using the high accuracy model and these reads were used for all subsequent analyses. Basecalled reads were mapped using Minimap2 (Li 2018) to the *L. rosalia* (NC_021952) and *L. chrysopygus* (NC_037878) mitogenomes. Resulting bam files were sorted and indexed using samtools and converted to bed files for viewing in IGV with bamToBed from bedtools (Li et al. 2009; Quinlan and Hall 2010). Visualization of read statistics was performed with Bamstats (http://bamstats.sourceforge.net).

### Mitogenome assembly

Assembly proceeded in two steps. We used the total reads aligning to *L. chrysopygus* mtDNA in fasta format to generate the assembly. Flye v2.8.3 was used for draft assembly and first round polishing of the initial consensus(Kolmogorov et al. 2020). Flye has an expected error rate of 0.5 to 1% for ONT reads and therefore was followed by an additional polishing step. Medaka v1.3.2 uses neural network models applied to a pileup of individual sequencing reads against a draft assembly to improve consensus sequences. In our testing, it outperformed Nanopolish in generating a contig without frameshift indels in the final contig (Simpson et al. 2017). Both packages were used with default settings.

### Circularization and annotation

Flye generated a circularized assembly, i.e. no repeats on the ends, ready for annotation. Identification of protein coding regions was performed through the MITOS2 website (http://mitos2.bioinf.uni-leipzig.de). Start position was manually set at the *COX1* gene following convention. The genome was visualized in Open Vector Editor (https://github.com/TeselaGen/openVectorEditor) and submitted to NCBI.

### DNA methylation analyses

Megalodon extracts high accuracy modified bases and variant calls from raw nanopore reads by using intermediate output from the neural network model provided by the basecaller Guppy. We used Megalodon v2.3.1 with the Guppy v4.6.4. The following research model from Rerio repository, res_dna_r941_min_modbases_5mC_5hmC_v001.cfg, was used as it is able to call both 5mC and 5hmC modifications at cytosine sites in any context. The “--mod-binary-threshold=0.8” flag was set to slightly decrease the stringency of Megalodon’s modification calling in line with suggested practices (Goldsmith et al. 2020). Further details are in supplementary file 2.

## Data Access

The mitogenome sequence has been deposited into NCBI GenBank with accession number MZ262294 and is available as supplementary file 3. Reads used to generate this assembly are available as supplementary file 4.

## Acknowledgements

We thank Executive Director Andria Heath, Director of Administrative Operations Nikki Sheehan and the keeper and animal care staff of the Utica Zoo as well as their tamarins Arie and Kane for generous donation of fecal material. We also thank A. Barks for experimental suggestions with DNA methylation analysis. We thank Sydney McGraw for the tamarin drawing. This work was supported by USDA AES Project. No. MIN-16-129, the UM Informatics Institute, and the SUNY Polytechnic Institute Research Foundation. The authors have no conflicts of interest and declare no competing financial interests.

## Author contributions

CF conceived and designed the study. AM obtained the samples. CF performed library prep and sequencing. NM, CF, and PL analyzed and interpreted the data. NM, AM, and CF drafted the manuscript. All authors reviewed and approved the manuscript.

## Abbreviations

CpG: Cytosine-phosphate-guanine
ONT: Oxford Nanopore Technologies
5mC: 5’methylation
5mCpG: 5’methylation at CpG site
5hmC: 5’hydroxymethylation
5hmCpG: 5’hydroxymethylation at CpG site
CH: Cytosine plus any non-G base

## Supplementary Data

Supplementary Figure S1. Alignment of reads to prior L. rosalia reference (NC_021952) vs. L. chrysopygus reference (NC_037878)

Supplementary File 1. Detailed methods and rationale.

Supplementary File 2. Table of methylation frequency by cytosine.

Supplementary File 3. Consensus sequence of *L. rosalia* mitogenome.

Supplementary File 4. Alignment reads used for mitogenome assembly.

